# libpspm: A feature-rich numerical package for solving physiologically structured population models

**DOI:** 10.1101/2023.08.04.551683

**Authors:** Jaideep Joshi, Lai Zhang, Elisa Z. Stefaniak, Ulf Dieckmann, Åke Brännström

## Abstract

For a vast majority of organisms, life-history processes depend on their physiological state, such as body size, as well as on their environment. Size-structured population models, or more generally, physiologically structured population models (PSPMs), have emerged as powerful tools for modelling the population dynamics of organisms, as they account for the dependences of growth, mortality, and fecundity rates on an organism’s physiological state and capture feedbacks between a population’s structure and its environment, including all types of density regulation. However, despite their widespread appeal across biological disciplines, few numerical packages exist for solving PSPMs in an accessible and computationally efficient way. The main reason for this is that PSPMs typically involve solving partial differential equations (PDEs), and no single numerical method works universally best, or even at all, for all PDEs. Here, we present libpspm, a general-purpose numerical library for solving user-defined PSPMs. libpspm provides eight different methods for solving the PDEs underlying PSPMs, including four semi-implicit solvers that can be used for solving stiff problems. Users can choose the desired method without changing the code specifying the PSPM. libpspm allows for predicting the dynamics of multiple physiologically structured or unstructured species, each of which can have its own distinct set of physiological states and demographic functions. By separating model definition from model solution, libpspm can make PSPM-based modelling accessible to non-specialists and thus promote the widespread adoption of PSPMs.

## Introduction

Organisms from the plant and animal kingdoms vary over 21 orders of magnitude in body mass (Jungers, 1986). Even when focusing on a single individual, growth-driven variation in body size can span several orders of magnitude between juvenile and adult life stages. Such ontogenetic variation naturally gives rise to strongly size-dependent life-history processes and size-asymmetric interactions among individuals. Individual rates of growth, mortality, and fecundity (collectively known as demographic rates) are not only varying with body size but are also modulated by the environment, which may include components that are independent of population structure (the non-feedback environment, e.g., temperature) and components that are affected by the population structure (the feedback environment, e.g., the abundance of a shared resource). Furthermore, the size dependence of life-history processes leads to size-dependent demographic trade-offs, i.e., trade-offs between growth, survival, and reproduction, which have led to the evolution of diverse life-history strategies. In addition to body size, usually measured in terms of length or weight, fundamentally different continuously variable dimensions of physiological population structure might be important too, such as gonad sizes, body conditions, energy reserves, metabolite concentrations, morphological shapes, symbiont densities, and microbiome compositions. Therefore, realistic predictions of the structure, dynamics, and life-history strategies of populations of organisms typically require accounting for a population’s physiological structure and its feedbacks with the environment.

Several modelling approaches exist for solving models with explicit physiological structure. In two of the most widespread and complementary approaches, population-level properties emerge from individual physiology and behaviour (Nisbet et al., 2016). They are: (1) individual-based models, also called agent-based models (ABMs), and (2) physiologically structured population models (PSPMs). ABMs are the most general approach to population modelling (Grimm, 1999). They are simpler to formulate, as they explicitly represent populations as collections of individuals and describe life-history processes and events at the level of individuals. However, ABMs are computationally intensive and, owing to the need for describing a finite number of agents, subject to demographic stochasticity. PSPMs (de Roos, 1997) offer an alternative, deterministic approach, which works well for large spatially homogeneous populations. A PSPM typically involves solving a transport equation, which is a partial differential equation (PDE) that describes the flow of individuals within a state space. Another approach for describing PSPMs based on renewal equations has also been developed (Diekmann et al., 2020), but it is better suited to model inter-generational change, whereas PDEs can describe the dynamics of a population in continuous time.

Specifying a PSPM using additional assumptions leads to three other classes of simpler models: (1) PDEs discretised in time, (2) PDEs discretised in state, and (3) PDEs discretised in both time and state. First, if the underlying PDE is discretized in time, we get a class of approaches called Integral Projection Models (IPMs), which can project a continuous state distribution from one time step to the next. The projection depends on demographic rates, which are typically estimated from observed data (Doak et al., 2021; Merow et al., 2014). Second, if we discretize the PDE only in state, we get stage-structured models, suitable for organisms with naturally discrete life-history stages. They entail solving a system of ordinary or delay differential equations (de Valpine et al., 2014). Finally, discretizing in both time and state leads to matrix population models (Crone et al., 2011), which are mathematically tractable and computationally efficient but are suitable only for organisms with naturally discrete life-history stages with constant probabilities of transitioning between stages. In principle, a finely state-discretized version of a PDE-based PSPM can be thought of as a matrix population model with time-varying matrix coefficients. In that sense, PSPMs combine the best of both worlds, i.e., the generality of ABMs with the computational tractability of matrix population models.

Several fundamentally different numerical schemes have been developed for solving transport equations in the context of fluid dynamics, meteorology, oceanography, and geophysics. However, standard PDE solvers that have been developed for such problems are not best suited for PSPMs because of the following reasons: (1) PSPMs have a solution-dependent boundary condition, (2) PSPM solutions have non-local dependencies, and (3) some problems are stiff (i.e., while solving them with explicit schemes, the numerical time integration breaks down or gets stuck with unreasonably small time steps), and require some amount of trial and error in choosing the best-suited numerical scheme for solving them. Existing software packages for solving PSPMs are limited in terms of the kinds of PSPMs that can be solved and the methods used to solve them. Examples include (1) EBTtool, which is the numerical solver in the PSPManalysis package (de Roos, 2021) and can solve PSPMs with multi-dimensional individual states and a distributed boundary condition but implements only one numerical method, the Escalator-Boxcar-Train (EBT), (2) Mizer (Scott et al., 2014), which can handle multiple species but only solves size-structured and thus one-dimensional models in the limited context of fisheries with a hard-coded biological model, and (3) Plant (Falster et al., 2011, 2017), which solves size-structured models using the characteristic method (CM) but only in the limited context of plant communities with a hard-coded biological model.

Wider adoption of PSPM modelling will be greatly facilitated by a computational tool that is (1) ecologically versatile by being applicable to a broad class of user-defined PSPMs, potentially with multiple interacting species, and (2) computationally flexible by allowing the application of multiple numerical methods to the same problem without having to change any model-specification code. Here, we present libpspm, a general-purpose numerical package for solving PSPMs. Compared to existing alternatives, the strength of libpspm lies in the following key features: (1) it enables users to define their own PSPM by specifying functions to compute the demographic rates and environmental conditions, (2) it provides users with the ability to switch, without making any modifications to the code, between eight different numerical methods, including semi-implicit methods for solving stiff problems, and (3) it allows users to model multispecies communities, with each species represented by either a structured or an unstructured population and with each species potentially having its own set of physiological states and demographic functions. By providing easy access to the numerical solution of the underlying PDE, libpspm will facilitate the widespread application of PSPMs to a variety of problems by separating ecological model definition from computational model solution, enabling researchers to focus on their system’s ecology.

## Methods

### PSPM definition

A physiologically structured population model (PSPM) describes how the distribution of individuals in the physiological state space evolves through time. The state space is spanned by *n* individual physiological state variables (*i*-state variables; *x*_*1*_, *x*_*2*_, …, *x*_*n*_).

At its core, a PDE-based PSPM is typically represented by the McKendrick-von Foerster equation,

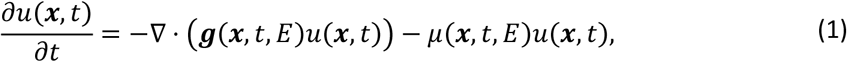

where vector quantities are expressed in bold letters; *x = [x*_*1*_, *x*_*2*_, …, *x*_*n*_]^*T*^ is the i-state variable, *u(x)* is the density of individuals in state space, such that the number of individuals within a small volume *dx* centred at *x* is *u(x)dx*, ***g***(.*), u(*.), and *β(*.) are the demographic rates of individual growth, mortality, and reproduction, respectively, as functions of the i-state *x*, the environment **E**, and time *t*. The growth rate ***g*** specifies the rate of change of all components of the i-state,

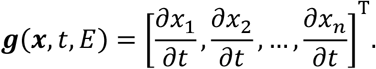

The first term on the right-hand side of Eq. 1 represents the change in the density of individuals in state *x* due to growth, whereas the second term represents the decrease in the density of individuals due to mortality.

Assuming that all individuals are identical at birth with i-state *x*_*b*_, Eq. 1 has an additional condition for the population density at the boundary,

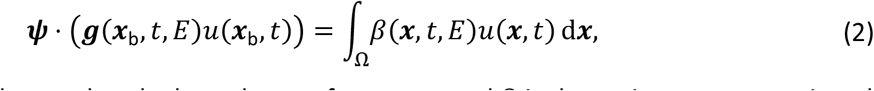

where *ψ* is the inward normal to the boundary surface at *x*_*b*_, and *Ω* is the entire state space, i.e., the domain of *x*.

Eq. 1 also has a specified initial condition,

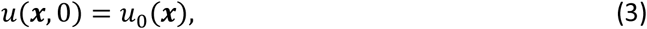

The boundary condition represents the fact that the inflow of individuals entering state *x*_*b*_ due to the birth of newborn individuals (produced by all individuals in the population; right-hand side of Eq. 2) must equal the outflow of individuals exiting state *x*_*b*_ due to the growth of newborn individuals (left-hand side of Eq. 2).

The feedback environment **E** is affected by all individuals in the population and thus often involves computing integrals over the state space. It typically takes the form

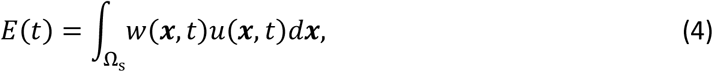

where *w* is some weighting function and *Ω*_s_ is some subset of *Ω*.

### PSPM definition for one-dimensional state space

PSPMs often consider only one state variable, typically the body size or related variables such as body weight and body energy reserves. For a one-dimensional state space, Eqs. 1-3 reduce to

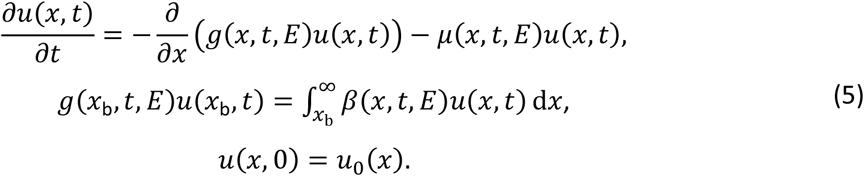

This work focuses on one-dimensional PSPMs with one or more species. However, libpspm will support multi-dimensional state spaces in the near future.

### Numerical solution

To solve the McKendrick-von Foerster equation, four numerical schemes are commonly employed: (1) the fixed-mesh upwind method (Sulsky, 1993; Ackleh & Ito, 1997; Fischer et al., 2006; Krzyzanowski et al., 2006; Hartvig et al., 2011), (2) the characteristic method (Angulo et al., 2014, 2016; Angulo & López-Marcos, 2002, 2004; Falster et al., 2016; Ito et al., 1991; Kostova, 2003), (3) the escalator-boxcar-train method (Brännström et al., 2013; de Roos, 1988), and (4) the agent-based method (Nisbet et al., 2016). We implement two versions of the first three schemes, one with an explicit integration method and one with a semi-implicit integration method. The agent-based method can be used as a baseline for comparison and as the main solver method for high-dimensional models. For the semi-implicit fixed-mesh upwind method, a first-order implementation is – as we show in the results – highly robust but prone to high levels of numerical diffusion. Therefore, we also provide an option to use a second-order version of this method, also called the linear upwind differencing method. Furthermore, we implement two state-of-the-art explicit ODE solvers from which users can choose – Runge-Kutta 4-5 Cash-Karp (Cash & Karp, 1990) and LSODA (Hindmarsh & Petzold, 2005). Thus, we have a total of eight solver methods: (1-2) explicit and semi-implicit fixed-mesh upwind methods (FMU and IFMU), (3) semi-implicit linear upwind differencing method (ILUD), (4-5) explicit and semi-implicit characteristic methods (CM and ICM), (6-7) explicit and semi-implicit EBT methods (EBT and IEBT), and (8) agent-based method (ABM). A conceptual description of the FMU, EBT, and CM methods can be found in (Zhang et al., 2017). A full description and implementation details of all the methods can be found in the Supplementary Information.

### Test models

To demonstrate the application of the aforementioned methods to PSPMs, we have chosen three reference models – the first two have analytical equilibrium solutions, allowing us to use them to assess the accuracy of the numerical methods. They also have specific modelling requirements, allowing us to showcase the features of libpspm. The third model is a complex vegetation model, allowing us to demonstrate not only the successful application of libpspm to a real-world problem, but also to construct and demonstrate certain problematic situations in which some numerical methods may break down or give divergent solutions.

1. The first model, from the animal kingdom, describes a size-structured population of a water flea, *Daphnia*, feeding on an unstructured algal resource (De Roos et al., 1990). The growth rate of the fleas depends on body size and algal resource availability, whereas the algae get consumed by the fleas and follow logistic regeneration dynamics. This model requires the solver to handle both structured and unstructured species and has the characteristic that the flea size distribution has a singularity at a certain size because fleas cease to grow beyond that size due to competition for food.
2. The second model, from the plant kingdom, is a simplified version of a vegetation demographics model called RED (for Robust Ecosystem Demographics), which represents a mass-structured population of trees (Argles et al., 2019). Each tree grows allometrically, and the recruitment of seedlings depends on the shading caused by standing trees. This model has the feature that plant size (measured in terms of individual plant biomass) can span six orders of magnitude, and the density distribution can reach extremely small values (∼*1*0^−*2*0^ individuals per unit area per unit size) at higher sizes.
3. The third model, again from the plant kingdom, is the ‘Plant’ model that describes a height-structured mixed-species community of plants competing for light (Falster et al., 2011, 2017). Taller plants shade other plants below them. Plant growth, mortality, and fecundity rates depend on their photosynthetic rates, which in turn depend on the light availability throughout their vertically extended crowns. The germination of seedlings also depends on the light availability near the ground. Competition between species leads to successional dynamics, such that early successional species are gradually replaced by mid and late successional species. In our implementation, species are characterized by their leaf mass per unit leaf area (LMA), which affects the investment in leaf biomass and leaf turnover rate, and consequently affects their demographic rates. The peculiarity of this model is that it can become stiff under certain conditions.

### Criteria for evaluation of numerical methods

We evaluate the performance of the numerical methods using the following criteria. (1) accuracy – this is defined in terms of the ‘biomass relative error’ *e* at equilibrium,

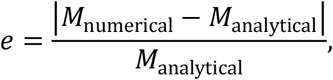

With

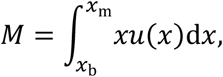

where *M* is a mathematical analogue of the total population biomass; (2) efficiency – this is measured in terms of execution time, and (3) robustness – this is a qualitative measure. A method is robust if it does not break (i.e., does not predict negative or exponentially increasing densities) and does not get stuck with unreasonably small timesteps. The former typically happens when mortality rates become too high, whereas the latter typically happens with stiff problems. We also qualitatively assess the methods based on the amount of numerical diffusion in the predictions, and the method’s versatility (e.g., whether it can be readily extended to multi-dimensional physiological states).

### Libpspm

libpspm is a numerical library for solving user-defined PSPMs. For best computational performance, it is implemented in C++, with an intuitive interface that allows defining, setting up, and simulating PSPMs in three easy steps (Box 1). Box 2 lists key features available for additional model specification and performance optimization. libpspm is open source and is available here: https://github.com/jaideep777/libpspm. Detailed package documentation and tutorials are also available at the associated package website: https://jaideep777.github.io/libpspm. Implementations of the test models and code to reproduce the figures in this paper is available in the ‘demo’ folder in the package.

#### Box 1.

Defining and solving a custom model with libpspm

##### libpspm – PSPM modelling in three steps

###### (1) To define a PSPM

define two classes for the Environment and the Individual (not necessarily with those names). These classes must inherit from the EnvironmentBase and IndividualBase classes provided by the library. In the derived classes, override the following member functions (one in Environment and four in Individual):

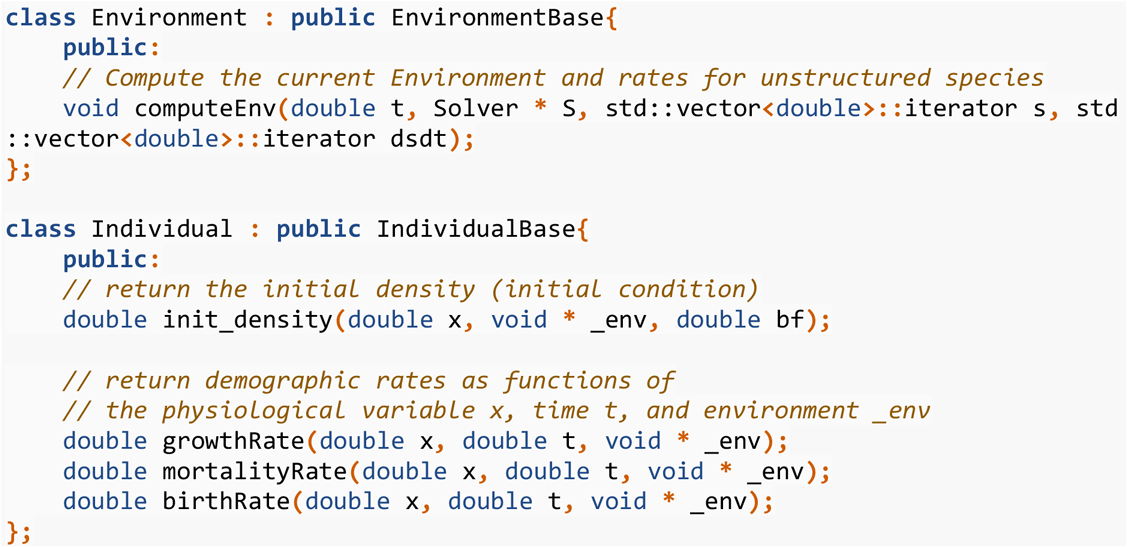

###### (2) To set up the PSPM solver

create an Environment object, create a species of Individuals, create a Solver by specifying the PSPM method and the ODE solver, and add the species and the environment to the solver. While creating the solver, it is possible to choose between the eight solver methods simply by changing the name of the method here, without any change to the rest of the code.

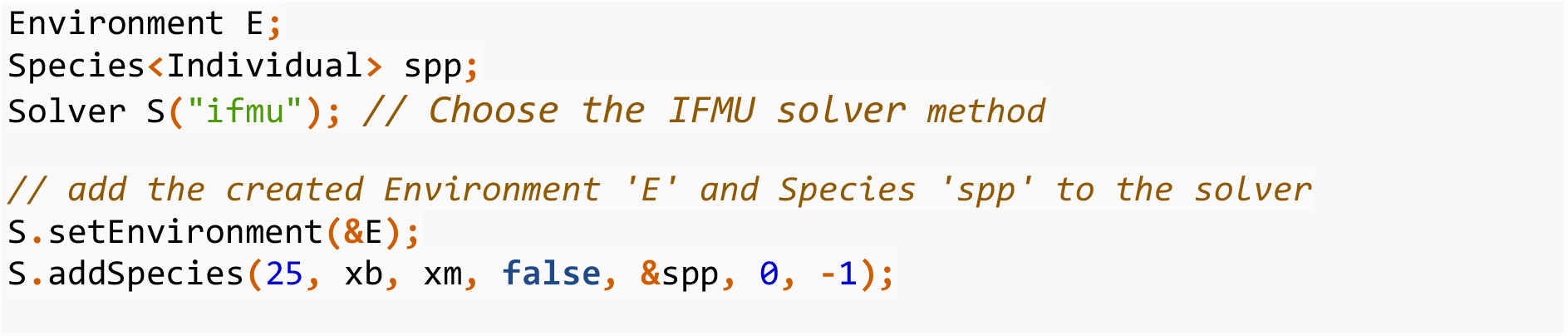

###### (3) To solve the PSPM

simply initialize and step the solver to the desired final time:

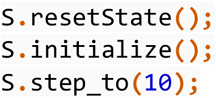

#### Box 2.

Additional libpspm features for specific modelling requirements and fine-tuning performance.

##### libpspm – Additional features

We briefly describe the additional features offered by libpspm for further model specification and performance optimization. Tutorials on their usage are available with the package.

###### (1) Computing integrals over state

State integrals are often required in the environment computation, but are computed differently for each discretization scheme. libpspm provides two functions to compute state integrals over the full range of states or above a threshold x_low, internally taking into account the method used by the solver.

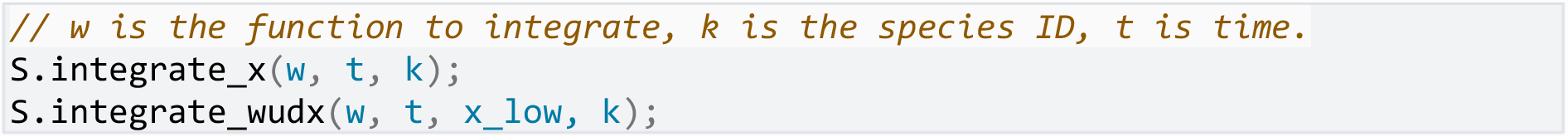

###### (2) Unstructured species

To create unstructured species, it is possible to add ‘system variables’ to the solver, which are also stepped using the internal ODE solver. Their rates can be computed in the same function defined for environment computation.

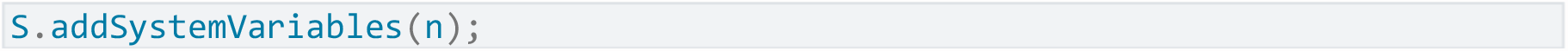

###### (3) Precomputation of the most intensive model components

Often, the demographic rates depend on a common quantity that is expensive to compute. E.g., in the Plant model, all rates depend on the photosynthetic rate. In such cases, the expensive quantity can be calculated in a separate function called ‘precompute’. The solver calls this function on-demand, i.e., a precompute is triggered by a change in the individual’s state or by a change in the environment, and the precomputation is executed just before the next demographic-rate computation.

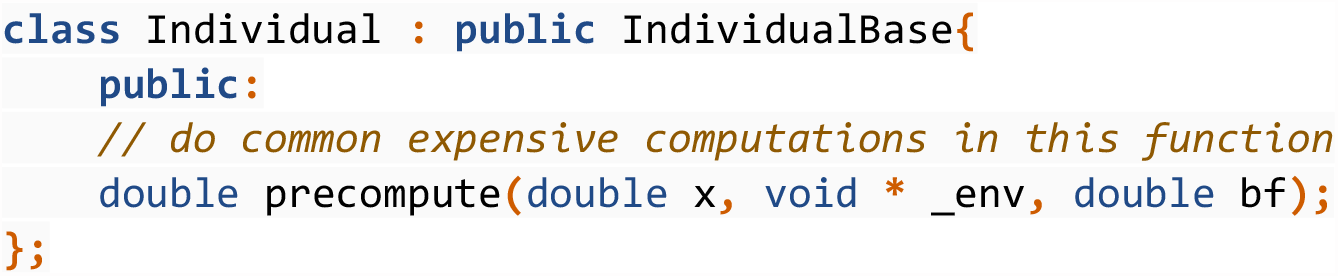

###### (4) Cumulative variables

It is often desirable to track some cumulative quantities for individuals, i.e., quantities integrated over time. For e.g., we might be interested to know the cumulative mortality of individuals, which is defined as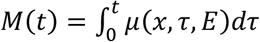. Such variables can be added while setting up the solver by defining the initialization and rate calculation for them. The solver then integrates them together with all other state variables.

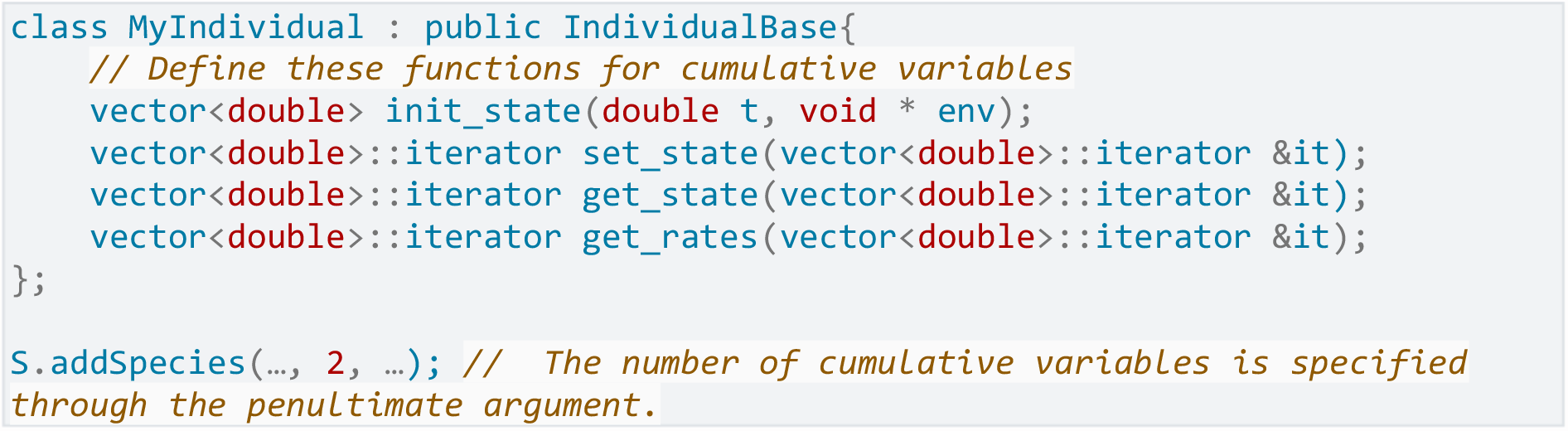

## Results

We first demonstrate the application of our eight numerical methods to the first two test problems – RED and Daphnia. We compare the analytical equilibrium size distributions with those simulated by libpspm. Among explicit methods (Fig. 1), all four are stable, implying that the RED and Daphnia models are not stiff. EBT and CM have the best accuracy for the RED model, whereas FMU has the best accuracy for the Daphnia model. CM and EBT end up with a lower resolution in the lower body-size range in the Daphnia model. At computationally reasonable population sizes, the ABM gives highly noisy solutions for the Daphnia model and is unable to capture higher size classes in the RED model. Among the semi-implicit methods (Fig. 2), all methods give good predictions of the equilibrium size distribution for the Daphnia model. However, the transient size distributions predicted by the IFMU and ILUD methods suffer from numerical diffusion and predict smoother size distributions than expected. In the RED model, the ILUD method breaks at higher sizes, where it predicts negative densities. The IFMU method correctly predicts densities at lower sizes but diverges from the analytical solution at higher sizes. At a modest resolution, all methods correctly predict the temporal dynamics and equilibrium values of population-level reproduction rates in both the RED and Daphnia models, in spite of numerical diffusion in the semi-implicit upwind methods (Fig. 3).

**Fig. 1.**
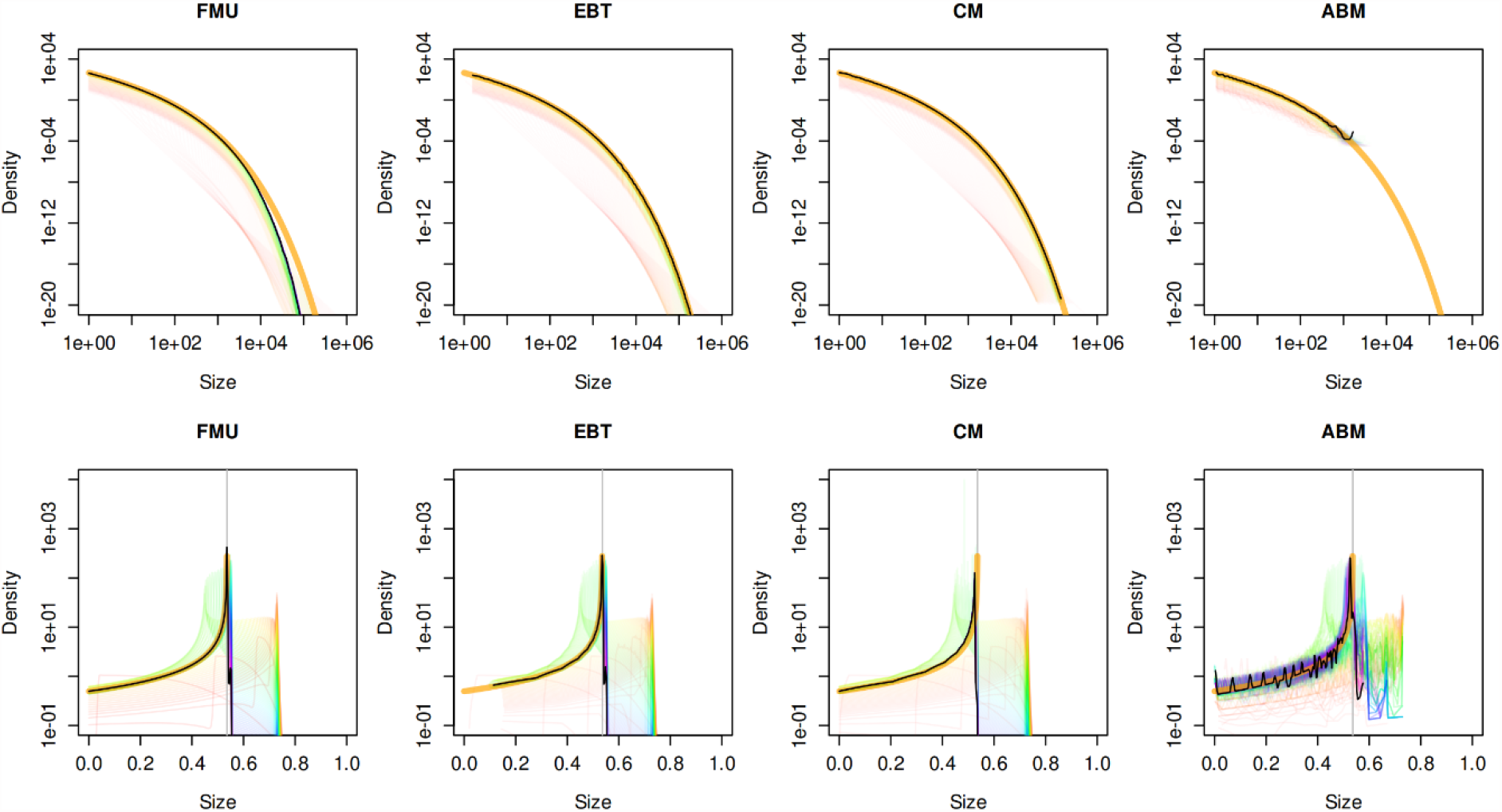
Explicit methods perform well for the RED and Daphnia models. Numerical equilibrium size distributions (black lines) for the RED (top row) and Daphnia (bottom row) models compared with analytical solutions (thick orange lines) for the four explicit methods, FMU, EBT, CM, and ABM. Thin coloured lines show transient solutions, going from red at *t =* 0 to purple for the last simulated time.

**Fig. 2.**
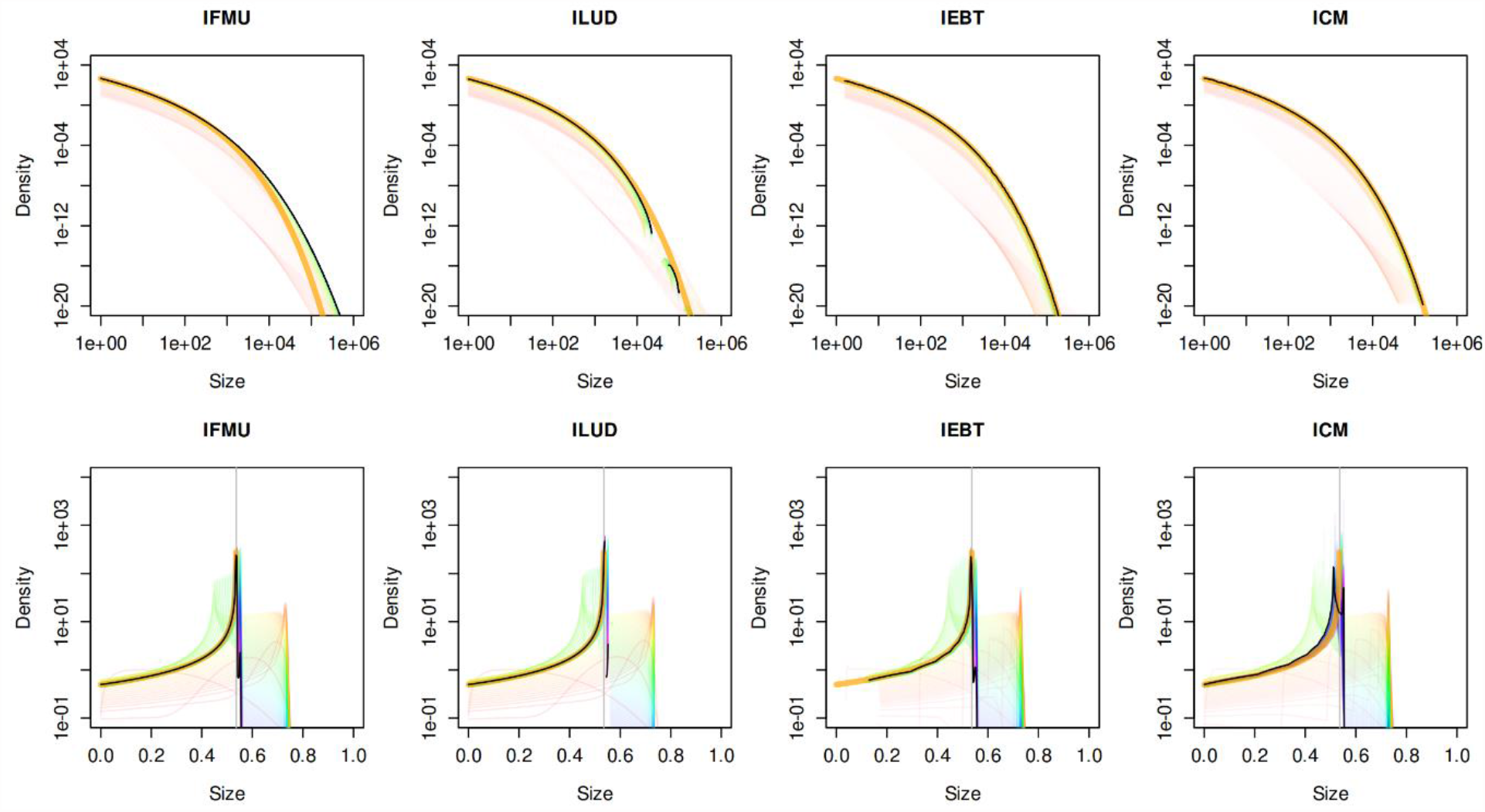
All implicit methods work well for the RED and Daphnia models. Numerical equilibrium size distributions (black lines) for the RED (top row) and Daphnia (bottom row) models match analytical solutions (thick orange lines) for the three semi-implicit methods – IFMU, ILUD, IEBT, and ICM. Thin coloured lines show transient solutions, going from red at *t =* 0 to purple for the last simulated time. Numerical diffusion can be clearly seen in the IFMU and ILUD methods, leading to smoother transient size distributions compared to the IEBT and explicit methods.

**Fig. 3.**
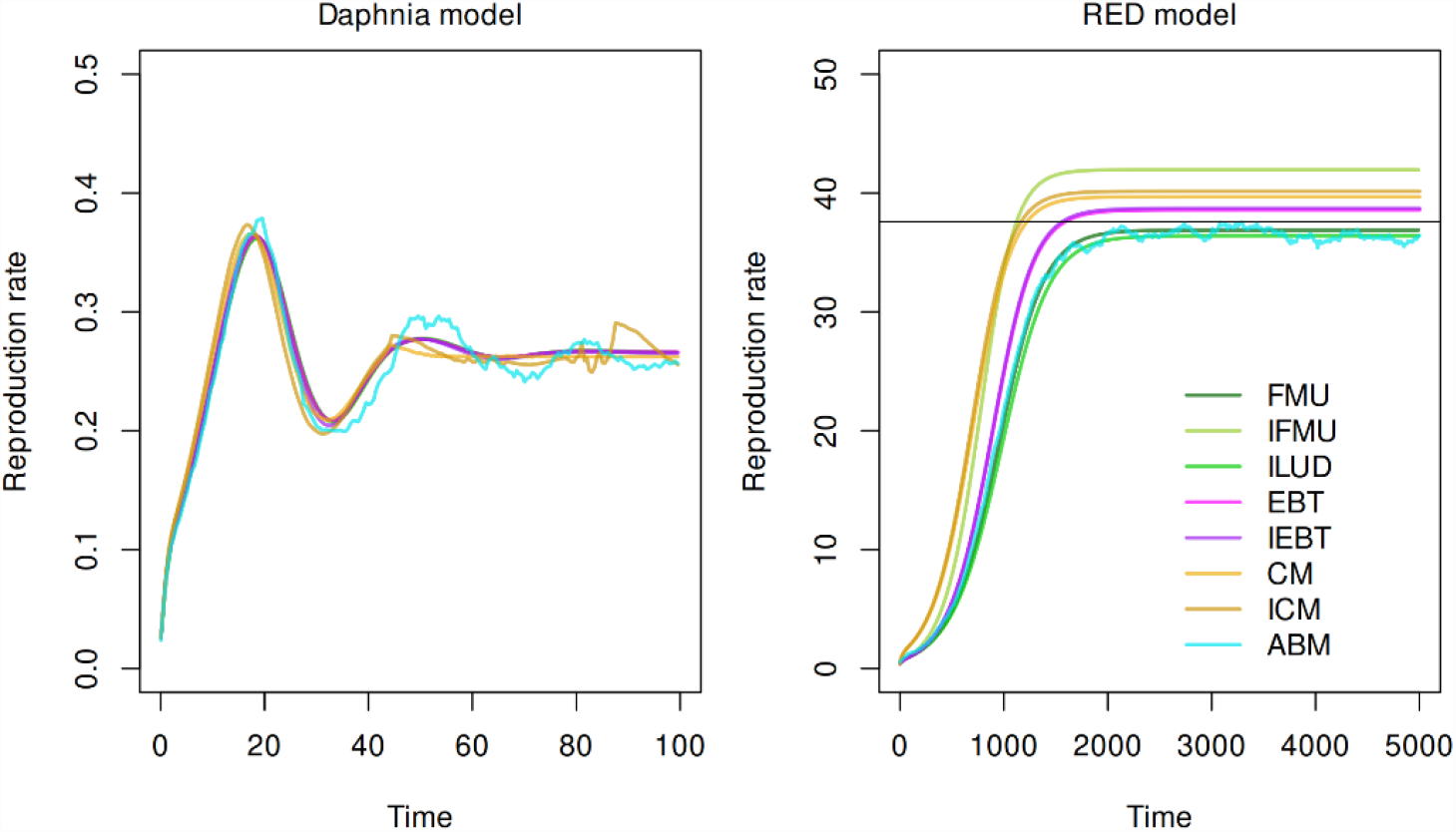
The eight methods differ slightly in their predictions of emergent population-level properties in the RED and Daphnia models. Predictions of transient and equilibrium population-level reproduction rates in the two models differ slightly among the eight methods, but roughly agree with the theoretical value.

We compare the performance of the eight methods using two metrics: (1) accuracy, measured in terms of the biomass error, and (2) efficiency, measured in terms of execution time (Fig. 4). For the Daphnia model, versions of the upwind scheme (FMU, IFMU, and ILUD) can be simulated at much lower resolutions compared to other methods and have a higher accuracy at lower resolutions. However, the EBT method, despite having lower accuracy at lower resolution, quickly overtakes the upwind methods and emerges as the fastest method when higher levels of accuracy are desired. For the RED model, the CM and EBT vastly outperform the upwind methods, with an order-of-magnitude higher accuracy for the same execution times. The CM method has the worst, almost unacceptable, performance for the Daphnia model, as was also reported by (Zhang et al., 2017). Surprisingly, however, it emerges as the best performing method for the RED model. The ABM performs moderately well for both models and has only a modest improvement in accuracy with increasing resolution. The performance of the semi-implicit methods saturates with increasing resolution for both models, as the timestep becomes the bigger limiting factor at higher resolutions.

**Fig. 4.**
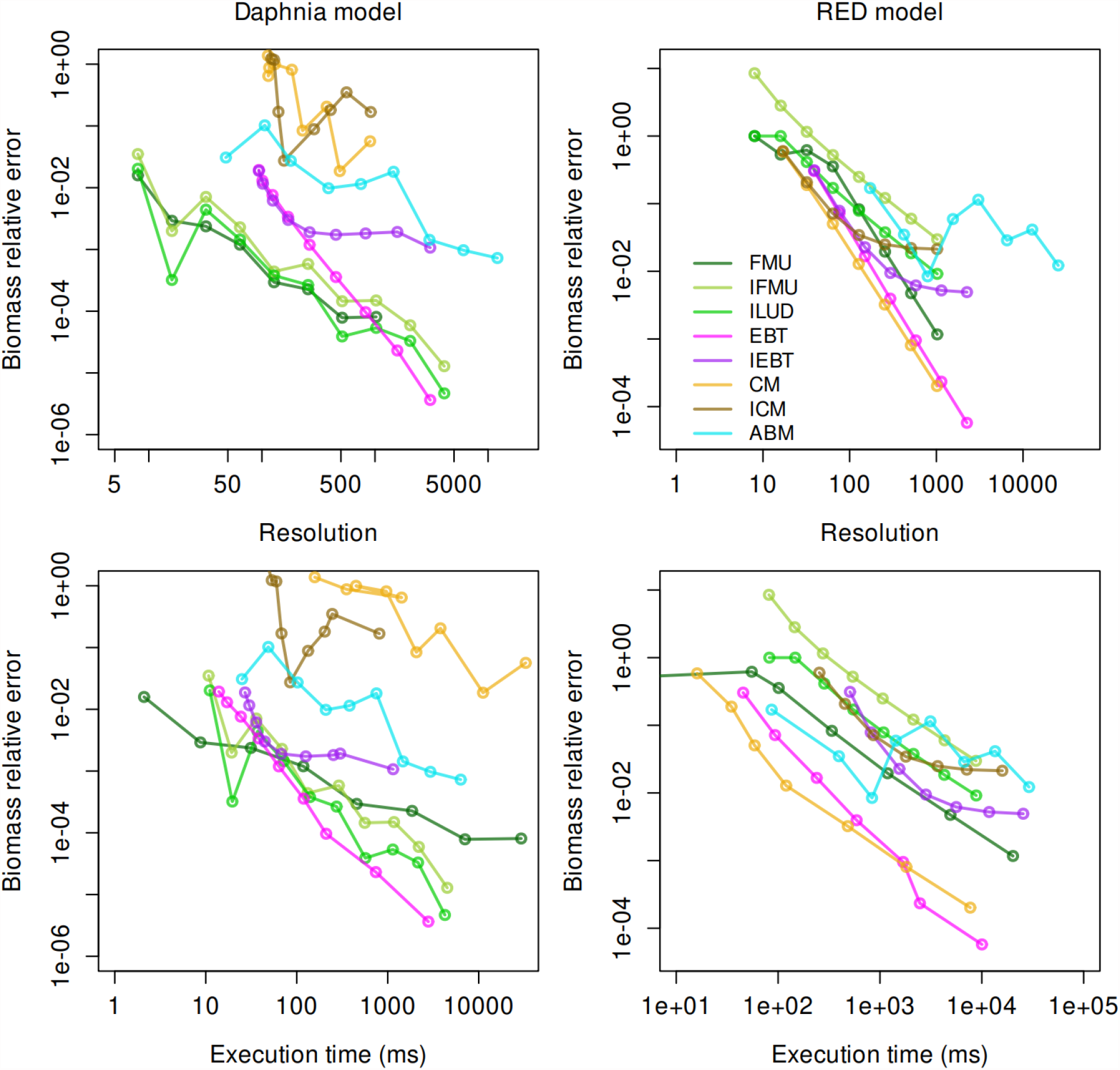
Performance ranking of the eight methods differs between models. Performance of the eight methods assessed in terms of the biomass relative error as a function of resolution (top row) and execution time (bottom row). Left panels represent the Daphnia model, whereas right panels represent the RED model.

Next, we apply the eight methods to ‘Plant’ – a complex multispecies vegetation model. This model can be simulated in two ways – (a) forced mode, in which the boundary condition is calculated with a fixed input ‘seed rain’ *I* and seed survival probability *s*_*e*_, such that *g*_*b*_*u*_*b*_ *= s*_*e*_*I*, i.e., the flux of offspring attempting to enter the population is held constant at each timestep, or (b) feedback mode, where the boundary condition is calculated as per Eq. 2. The forced mode is useful for predicting the equilibrium properties of a metapopulation of patches (Falster et al., 2017), whereas the feedback mode is useful for simulating the transient dynamics of a single patch. We simulated the model with three species, each with a different value of leaf mass per area (LMA). In the Plant model, species with low LMA are expected to be early successional and are thus expected to be replaced by species with progressively higher LMA. We compare the methods in terms of their predicted transient and equilibrium size distributions and reproductive output (seed rain). In forced mode, all eight methods predict similar transient and equilibrium solutions, albeit with varying resolutions (Fig. 6). The IFMU and ILUD methods predict incorrect transient seed rains at a resolution comparable to the other upwind methods but give the correct transient solutions when simulated with higher resolutions. This is because the IFMU has the highest numerical diffusion, as is also evident in its predictions of the size-distribution (Fig. 5), whereas the ILUD has the tendency to predict negative densities, especially if the timestep is not small enough.

**Fig. 5.**
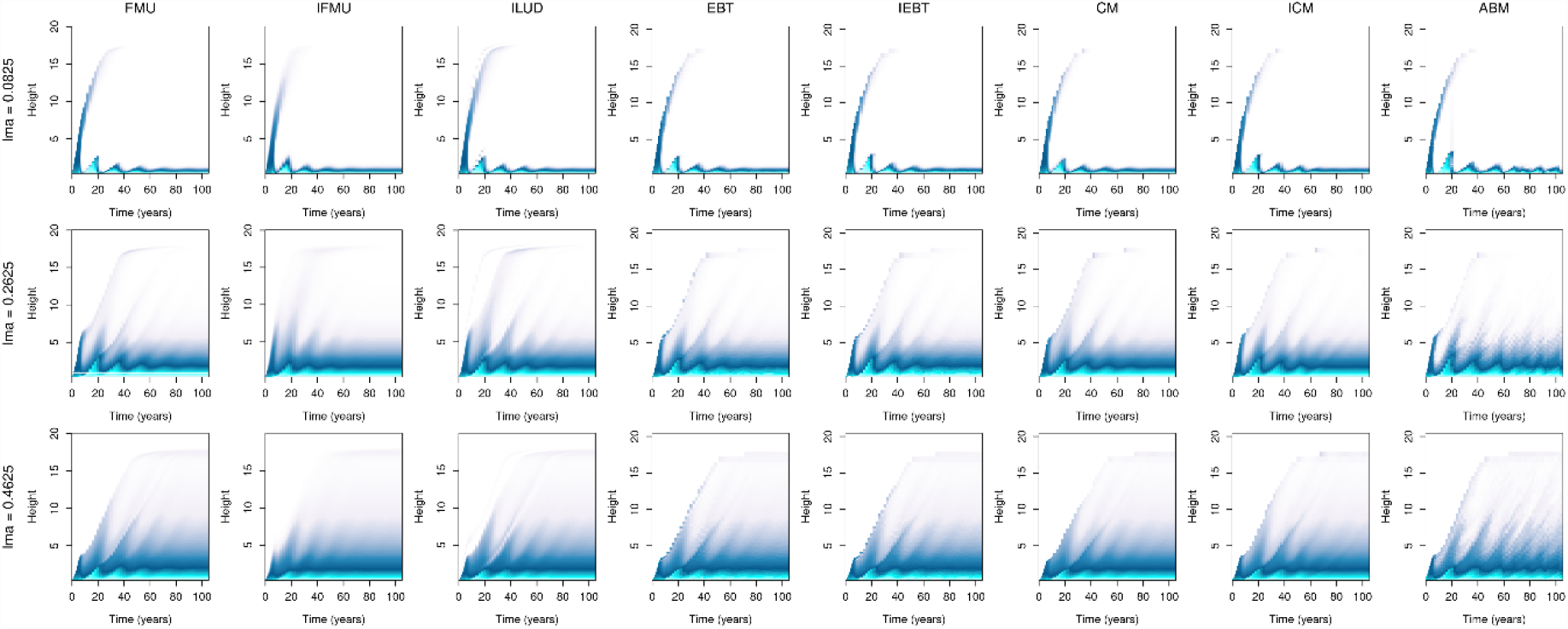
All methods correctly simulate the time-evolution of the size distribution of three species in the Plant model. For the Plant model simulated with a fixed input seed rain with three species with different LMAs, all methods broadly agree on the solutions.

**Fig. 6.**
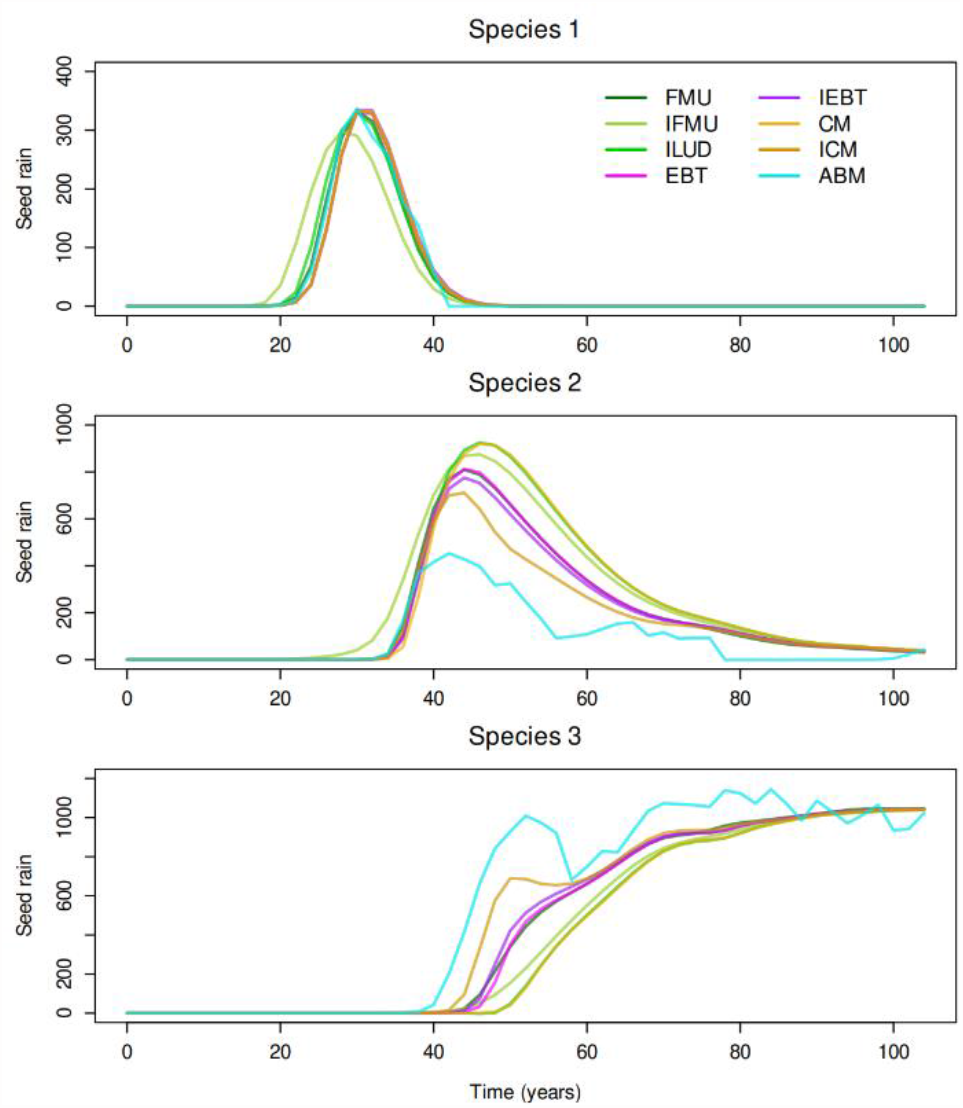
The eight methods differ slightly in their predictions of emergent population-level quantities in the Plant model. Timeseries of the reproductive output (seed rain) of three species in the Plant model forced with a fixed input seed rain. All methods mostly agree on the solution. The fixed-mesh methods did not give the same results at comparable resolution. Here the FMU method was simulated with 199 cells, ILUD was simulated with 499 cells, and IFMU with 1069 cells. At lower resolution, the ILUD method predicts negative seed rains for parts of the trajectory, and the IFMU solver predicts seed rains with high errors.

The Plant model in feedback mode can become stiff, allowing us to construct and demonstrate two kinds of problematic situations one should be wary of in simulating real-world PSPMs. First, when starting from ‘bare ground’, with a minimal number of seeds, i.e., *u(x*, 0*) = u*_0_*δ(x* − *x*_*b*_), where *δ(x)* is the Dirac delta function, the eight methods do not agree on the transient solution. This is because the model is started with an extremely specific and rather artificial initial condition, leading to an accumulation of errors that are characteristic to the method and have little opportunity to average out. Thus, even slight differences in the initial condition (arising, for e.g., from the differences in discretization) lead to slightly different seed outputs, which feed back into the population and lead to divergent transients among the eight methods (Fig. S1). Indeed, this is a tricky initial condition that few numerical schemes are developed to handle. The solution to this problem is to ensure that the initial condition is slightly spread out. Second, when starting from an initial size distribution of the form *u(x*, 0*) = u*_0_*e*^−*x*/4^, the problem is stiff. Six of the eight methods break down in this case. The solution to this problem is to use semi-implicit methods: particularly, we found that the IFMU and IEBT remain stable (Fig. S2, Fig. S3). We also note that in this case, since the initial condition is more spread out, accumulation of errors is less characteristic of the discretization scheme, and the two methods agree on the transient solution (Fig. S2).

## Discussion

We have demonstrated the applicability and limitations of eight methods for solving PSPMs, and presented a numerical library that allows for simulating any user-defined PSPM with any of the eight methods. Broadly, the eight methods can be categorized in two ways – (1) grid-based (upwind methods) vs cohort-based (EBT and CM), or (2) explicit vs implicit. The advantage of cohort-based methods is that they are fast and highly accurate, but the disadvantage is that their predictions of the full density distribution suffer from low resolution at the lower end of the physiological axis. Among the explicit methods, we found that the EBT method consistently delivered the best performance and was the fastest method when high accuracy was desired. The IFMU method was the most robust and worked for all test problems, while the explicit methods (FMU, EBT, CM), ILUD and ICM did not work for the Plant model running in feedback mode. While the IEBT also worked in all situations, and should work for most practical problems, it is possible to conceive specific situations under which it could become unstable, for e.g., when the boundary cohort has a very high growth rate gradient. The IFMU method had the highest numerical diffusion, but this did not necessarily compromise its predictions of integrated population-level properties.

While the ABM’s performance was (unsurprisingly) significantly lower than ODE based methods, it is extremely versatile, and allows for modelling features that are difficult to incorporate in ODE-based methods. We illustrate this point with two examples. First, if the population is structured along multiple variables, i.e., the individual state has dimensionality *m* > *1*, the complexity of grid-based methods will increase exponentially with *m*, i.e., *O(n*^*m*^), where *n* is the grid resolution. On the other hand, the EBT and CM are more amenable to handle multi-dimensional state variables, because they can skip simulating empty regions of the state-space. The ABM is even more amenable to such situations, and can readily handle states of any dimensionality. Second, cohorts or grid-cells in the ODE-based methods consist of necessarily identical individuals. Although trait variation can be accounted for by simulating multiple ‘species’ differing in their trait values (as in the Plant model), the number of species scales exponentially with the number of traits required. The ABM can readily account for trait variation among individuals and species, but at the expense of demographic stochasticity that increases with the number of traits and state variables. Table 1 shows a comparison of all the methods with respect to five evaluation criteria.

**Table 1.** Comparison of the eight methods with respect to different criteria.

| Method | Accuracy | Efficiency | Robustness | Diffusion | Versatility |
| --- | --- | --- | --- | --- | --- |
| FMU | Moderate | Moderate | Moderate | Moderate | Low |
| IFMU | Low-moderate | Low-moderate | Guaranteed | Very high | Low |
| ILUD | Moderate | High | Low | Moderate | Low |
| EBT | Very high | Very high | Moderate | Zero | High |
| IEBT | Moderate | Very high | High | Zero | High |
| CM | Inconsistent | High | Moderate | Low | Moderate |
| ICM | Inconsistent | High | Moderate | Low | Moderate |
| ABM | Low | Very low | Guaranteed | Zero | Very high |

The question then arises, which numerical method is suitable for a given problem? In general, in the interest of accuracy, we recommend using the EBT or CM methods first, especially if only integrated population-level quantities are desired in the final outputs. If accurate predictions of the full density distribution are required, then we recommend using the FMU method. Some problems can be stiff. It is difficult to assess beforehand whether a specific problem is stiff or not. The best option is to try an explicit method, and if it breaks (that is an indication that the problem is stiff), the IFMU or IEBT methods are a good choice. Note that the CM method implemented here solves for point mass density, but it is possible to conceive a finite-volume implementation of this method, which could resolve the problem of non-uniform resolution across the range of states. Our recommendations are summarized in Table 2.

**Table 2.** Recommended methods in different situations.

| Method | Problem is not stiff | Problem is stiff |
| --- | --- | --- |
| Full density distribution desired | FMU | IFMU |
| Only integrated population-level quantities desired | EBT or CM | IEBT |

With advancements in computational infrastructure, biological models, aiming at incorporating greater realism, are continuously getting ever more complex. Incorporating the size structure of populations is one such dimension of complexity that has particularly come into focus, for e.g., in vegetation modelling with the advent of vegetation demographic models (VDMs). However, many such models, being already computationally intensive, do not allow for a rigorous testing of the numerical methods used to solve them. For example, some cohort-based vegetation models use a discretization scheme similar to the EBT but ignore the boundary cohort dynamics. We found that doing so introduces large errors in the numerical solutions (not shown). Thus, by providing multiple choices for handling computational complexity and separating model specification from numerical model solution, libpspm has the potential to play a critical role in development of the next-generation size-structured models.

## Acknowledgements

We acknowledge Mazouth-Laurol Maxime for initial work on a prototypical MATLAB-based package that inspired the development of libpspm. JJ and UD acknowledge funding from the European Union’s Horizon 2020 research and innovation programme under the Marie Skłodowska-Curie Actions fellowship (grant agreement No. 841283 – project Plant-FATE, Assessing the global vulnerability of forests to drought using plant functional trait evolution). JJ and EZS acknowledge funding from the Strategic Initiatives Program of the International Institute for Applied Systems Analysis (project RESIST, Resilience of ecosystem services provided by intact and sustainably managed terrestrial ecosystems). JJ, EZS, and UD acknowledge funding from the European Union’s Horizon 2020 research and innovation programme (grant agreement No. 820989 – project COMFORT, Our common future ocean in the Earth system: quantifying coupled cycles of carbon, oxygen, and nutrients for determining and achieving safe operating spaces with respect to tipping points; the work reflects only the authors’ view, and the European Commission and their executive agency are not responsible for any use that may be made of the information the work contains). UD acknowledges funding from the Japanese Society for the Promotion of Science, through a KAKENHI Start-up grant (Identifying tipping points and safe operating spaces in sustainable fisheries management under future climate change) and a KAKENHI C grant (Accounting for evolutionary and socioeconomic impacts in modern fisheries science and management). UD acknowledges funding from the Global Bioconvergence Center of Innovation at the Okinawa Institute of Science and Technology Graduate University, OIST, supported by a grant from the Japan Science and Technology Agency, JST, Program for an Open Innovation Platform for Academia-Industry Co-Creation, COI-NEXT. JJ, EZS, UD, and ÅB acknowledge funding from the National Member Organizations that support IIASA.

## Author contributions

JJ and ÅB designed the study with inputs from LZ and UD. JJ wrote the code, performed the analysis, and wrote the first draft of the manuscript. EZS contributed complementary code. All authors contributed to conceptual development and revision of the manuscript.

## Notes

https://github.com/jaideep777/libpspm

### Competing Interest Statement

The authors have declared no competing interest.

