## Supplementary Text and Figures for "libpspm: A feature-rich numerical package for solving physiologically structured population models"

Jaideep Joshi et al.

#### Supplementary text

Below, we first describe the test models, followed by a description of the eight methods to solve PSPMs.

##### Description of test models

Here, we briefly describe the implementation of the RED and Daphnia models. The 'Plant' model is implemented exactly as per (Falster et al., 2017), so we do not repeat the model description here.

###### RED model

Our implementation of the RED model is a simplified version of (Argles et al., 2019). The state variable $x$ represents individual plant biomass ($m$) normalized by a reference value of $m_0 = 1$ kg ($x = m/m_0$). Plant growth rate is allometrically related to their biomass,

$$g(x) = g_0 x^{\phi_{\mathrm{g}}}.$$

The mortality rate of plants is assumed constant,

$$\mu(x) = \mu_0.$$

Reproduction rate depends on the fraction $\alpha$ of energy invested into reproduction (vs growth) and the proportion of shaded area $E$,

$$\beta(x) = \frac{\alpha}{1-\alpha}\left(g_0 x^{\phi_{\mathrm{g}}}\right)(1-E).$$

Shaded area $E$, expressed in ha, in turn, depends on the total crown area $A_{\mathrm{c}}$ of all trees in the population. Assuming non-overlapping crowns,

$$E = E_0 \int_0^{\infty} A_{\mathrm{c}}(x) u(x) \mathrm{d}x.$$

Furthermore, crown area is also assumed to scale allometrically with plant biomass,

$$A_{\mathrm{c}} = a_0 x^{\phi_{\mathrm{a}}}.$$

The equilibrium density distribution for this model can be calculated as

$$u^*(x) = n_{\mathrm{eq}}\left(\frac{1}{E_0}\right)\left(\frac{\mu_0}{g_0}\right) x^{-\phi_{\mathrm{g}}} \exp\left(\chi\left(1 - x^{1-\phi_{\mathrm{g}}}\right)\right),$$

where

$$n_{\mathrm{eq}} = \frac{1}{a_0} \frac{\left(1 - \left(\frac{1-\alpha}{\alpha}\right)\left(\frac{(\mu_0/g_0)}{\chi^{\Phi_{\mathrm{g}}} \exp(\chi)\, \Gamma(\chi, \Phi_{\mathrm{g}}+1)}\right)\right)}{\left(\chi^{\Phi_{\mathrm{a}}} \exp(\chi)\, \Gamma(\chi, \Phi_{\mathrm{a}}+1)\right)},$$

where $\Gamma(a, x)$ is the upper incomplete gamma function and

$$\chi = \frac{(\mu_0/g_0)}{1-\phi_{\mathrm{g}}}, \qquad \Phi_{\mathrm{g}} = \frac{\phi_{\mathrm{g}}}{\phi_{\mathrm{g}}-1}, \qquad \Phi_{\mathrm{a}} = \frac{\phi_{\mathrm{a}}}{\phi_{\mathrm{g}}-1}.$$

Parameter values used in our simulation are given in table below.

| **Parameter** | **Symbol** | **Value** | **Unit** |
|---|---|---|---|
| Growth rate scalar | $g_0$ | 0.0838 | $\mathrm{year}^{-1}$ |
| Growth rate exponent | $\phi_{\mathrm{g}}$ | 0.7134 | - |
| Mortality rate | $\mu_0$ | 0.035 | $\mathrm{Year}^{-1}$ |
| Fraction of allocation to reproduction | $\alpha$ | 0.1 | - |
| Crown area scalar | $a_0$ | 0.396 | $\mathrm{m}^2$ |
| Crown area exponent | $\phi_{\mathrm{a}}$ | 0.749 | - |
| Environment scalar | $E_0$ | $10^{-4}$ | $\mathrm{m}^{-2}$ |

#### Daphnia model

Following (Zhang et al., 2017), we use the simplified version of the Daphnia model as introduced by (de Roos, 1988). Growth rate follows a Holling Type II functional response to food availability $S$, and growth is stunted if insufficient food is available,

$$g(x, t, S) = \max\left(\frac{S(t)}{1+S(t)} - x, 0\right).$$

The mortality rate is constant, giving

$$\mu(x,t,S) = \mu_0.$$

The reproduction rate is proportional to the energy acquired from food intake and increases with body size,

$$\beta(x,t,S) = \alpha x^2 \frac{S(t)}{1+S(t)}.$$

The resource follows logistic growth dynamics and gets depleted when consumed by daphnia,

$$\frac{\mathrm{d}S}{\mathrm{d}t} = rS\left(1 - \frac{S}{K}\right) - \frac{S}{1+S}\int_0^1 x^2 u(x,t)\mathrm{d}x.$$

This model has an analytical equilibrium solution,

$$u^*(x) = \frac{\alpha r S^* \left(1 - \frac{S^*}{K}\right)}{(x^*)^\mu}(x^* - x)^{\mu-1}, \qquad 0 \le x \le x^*,$$

with

$$S^* = \frac{x^*}{1-x^*}, \qquad x^* = \left(\frac{\mu(1+\mu)(2+\mu)}{2\alpha}\right)^{1/3}.$$

Parameters used in our simulations are given in the table below.

| **Parameter** | **Symbol** | **Value** | **Unit** |
|---|---|---|---|
| Scaled reproduction rate | $\alpha$ | 0.75 | - |
| Mortality rate | $\mu_0$ | 0.1 | day$^{-1}$ |
| Intrinsic growth rate of resource | $r$ | 0.5 | day$^{-1}$ |
| Carrying capacity of resource | $K$ | 3 | cells/dm$^3$ |

#### Description of numerical methods implemented in libpspm

Below, we briefly describe the implementation of the eight numerical schemes in libpspm. We do not delve into the conceptual details of the schemes, for which we refer the reader to relevant literature.

##### Explicit fixed-mesh upwind method (FMU)

The fixed-mesh solver relies on to discretizing the McKendrick von-Foerster PDE (Eq. 1) on a fixed grid (mesh) with $J$ cells. The edges of the gridcells are at $x_j, j \in [0, J]$, and centres are at $X_j = (x_j + x_{j+1})/2, j \in [0, J-1]$. Note that $x_0 = x_b, x_J \ge x_m$. The FMU method solves for the solution $u(x)$ at the cell centres $X_j$.

Integrating the PDE over a grid cell $j$, we get

$$\frac{\partial}{\partial t}\int_{x_j}^{x_{j+1}} u(x,t)\mathrm{d}x = -[g(x,t,E)u(x,t)]_{x_j}^{x_{j+1}} - \int_{x_j}^{x_{j+1}} \mu(x,t,E)u(x,t)\mathrm{d}x. \qquad \text{(S1)}$$

Let us define the average value of $u(x)$ over the cell $j$ as $U_j$. Therefore,

$$h_j U_j = \int_{x_j}^{x_{j+1}} u\,(x,t)\mathrm{d}x,$$

where $h_j$ is the width of the cell, $h_j = x_{j+1} - x_j$. We also approximate the integral on the RHS and the boundary condition using the midpoint quadrature rule,

$$\int_{x_j}^{x_{j+1}} \mu(x,t,E)u(x,t)\mathrm{d}x = h_j\mu(X_j)U_j,$$

$$\int_{x_b}^{x_m} \beta(x,t,E)u(x)\mathrm{d}x = \int_{x_0}^{x_J} \beta(x,t,E)u(x,t)\mathrm{d}x = \sum_{j=0}^{J-1} h_j\,\beta(X_j)U_j.$$

Therefore, we can write

$$\frac{\partial}{\partial t}U_j = -\frac{[g(x,t,E)u(x,t)]_{x_j}^{x_{j+1}}}{h_j} - \mu(X_j)U_j, \quad \text{(S2)}$$

which gives us the equation for calculating the required derivatives. For the explicit method, we integrate these derivatives using the Runge-Kutta 4-5 Cash-Karp integrator.

To calculate the first term on the RHS of Eq. (S2, we still need a way to calculate the density function at the cell edges, i.e., $u_j$. One conceivable way to do so is to simply interpolate it from the values of $U$ from the neighbouring cells. However, this solution is unstable.

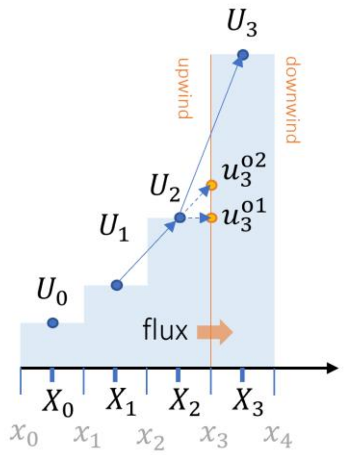

**Fig. M1. Calculating $u$ at cell boundaries in the FMU method.**

**The upwind scheme.** A more stable way to get the density at the cell edges is to use the upwind difference scheme. In the first-order upwind scheme, the value of $u$ is assumed to be constant throughout the grid cell, thus giving

$$u_3^{o1} = U_2.$$

While this method is more stable, it has low accuracy. The second-order upwind scheme assumes linear variation in $u$ within a grid cell. Thus, we can again find $u_3$ by linear interpolation, but this time, extrapolating it from the upwind values of $U$,

$$u_3^{o2} = U_2 + \Delta U(x_3 - X_2),$$

where

$$\Delta U = \frac{(U_2 - U_1)}{(X_2 - X_1)}.$$

This method is accurate, but again, can become unstable if the slope $\Delta U$ is too high. To obtain the best of both worlds, i.e., the stability of the first-order method and the accuracy of the second-order method, we can combine the two solutions as follows,

$$u_i = U_{i-1} + \phi \frac{(U_{i-1} - U_{i-2})}{(X_{i-1} - X_{i-2})}(x_i - X_{i-1}),$$

where $\phi$ is a scalar between 0 and 1. When $\phi = 0$, $u_i$ is obtained from the first-order scheme, and when $\phi = 1$, from the second-order scheme.

**Flux limiter.** The value of $\phi$ can be obtained as a function of the ratio $r$ of successive slopes of the density function,

$$r = \frac{\Delta U_{\text{downwind}}}{\Delta U_{\text{upwind}}},$$

where, e.g.,

$$\Delta U_{\text{downwind}} = \frac{U_i - U_{i-1}}{X_i - X_{i-1}}$$

and

$$\Delta U_{\text{upwind}} = \frac{U_{i-1} - U_{i-2}}{X_{i-1} - X_{i-2}}.$$

When the upwind slope is too high compared to the downwind slope ($r \to 0$), the second-order scheme will give a very high (undesirable) value of $u_3$. In such a case, we want the first-order scheme, and thus $\phi \approx 0$. When the upwind slope is too small compared to the downwind slope, the second-order approximation of $u_3$ will be too close to the first-order approximation, and we can improve accuracy by pushing it up even more ($\phi > 1$). All these features are achieved by the Superbee flux limiter,

$$\phi(r) = \max\Big(\max\big(0, \min(2r, 1)\big), \min(r, 2)\Big)$$

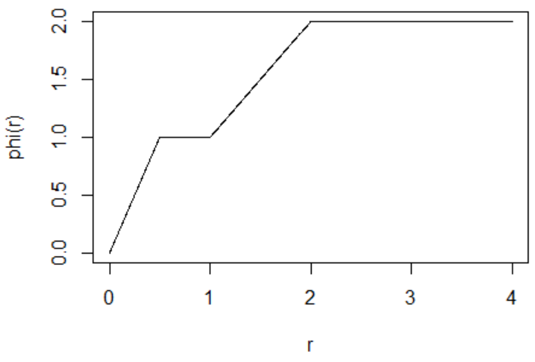

**Fig. M2. Superbee flux limiter.**

**Calculating $u_1$, $u_{J-1}$, and $u_J$.** Finally, we also need to find the values of $u_1$, $u_{J-1}$, and $u_J$, which cannot be found by the above formula due to out-of-bounds subscripts. We simply use the first-order scheme to get these,

$$u_i = U_{i-1}.$$

**Boundary condition.** $u_0$ is obtained from the boundary condition

$$u_0 = \frac{s_\mathrm{e}}{g(x_0)} \sum_{j=0}^{J-1} h_j\, \beta(X_j) U_j.$$

#### Semi-implicit first-order upwind method (IFMU)

The implicit versions of the fixed-mesh upwind scheme also discretize the state variable on a fixed mesh, but make the time update implicit (Hartvig et al., 2011). The points at which $u$ is evaluated can be specified to be the centres or lower edges of the grid-cells. The first-order (IFMU) and second-order (ILUD) methods rely on a first-order or second-order formula for calculating the derivative with respect to the physiological variable. Thus, the discretized version of Eq. 1 for the IFMU method is

$$\frac{u_j^{n+1} - u_j^n}{\Delta t} = \frac{g_j^n u_j^{n+1} - g_{j-1}^n u_{j-1}^{n+1}}{\Delta h_j} - \mu_j^n u_j^{n+1}. \quad \text{(S3)}$$

Note that these equations assume a semi-implicit time update, such that the demographic rates $g$ and $\mu$ are evaluated using the current state (index $n$), whereas the densities are evaluated at the next state (index $n+1$). A fully implicit method would evaluate all quantities on the RHS at the next state, but this is not computationally feasible as it would involve high-dimensional root-finding.

Equation (S3 can be used to find successive values of $u_j^{n+1}$ as follows,

$$\begin{aligned} u_0^{n+1} &= B/g_0^n, \\ u_j^{n+1} &= \frac{u_j^n + u_{j-1}^{n+1} g_{j-1}^n \dfrac{\Delta t}{\Delta h_j}}{1 + \mu_j^n \Delta t + g_j^n \dfrac{\Delta t}{\Delta h_j}}, \\ B &= s_\mathrm{e} \sum_{j=0}^{J-1} \beta_j^n u_j^n h_j. \end{aligned} \quad \text{(S4)}$$

#### Semi-implicit first-order upwind method (ILUD)

The discretized version of Eq. 1 for the ILUD method is

$$\frac{u_j^{n+1} - u_j^n}{\Delta t} = \frac{3 g_j^n u_j^{n+1} - 4 g_{j-1}^n u_{j-1}^{n+1} + g_{j-2}^n u_{j-2}^{n+1}}{2\Delta h_j} - \mu_j^n u_j^{n+1}. \quad \text{S5}$$

The successive values of $u_j^{n+1}$ can then be calculated as follows,

$$
\begin{aligned}
u_0^{n+1} &= \frac{B}{g_0^n},\\
u_1^{n+1} &= \frac{u_1^n + u_0^{n+1} g_0^n \frac{\Delta t}{\Delta h_j}}{1 + \mu_0^n \Delta t + g_1^n \frac{\Delta t}{\Delta h_j}},\\
u_j^{n+1} &= \frac{u_j^n + u_{j-1}^{n+1} g_{j-1}^n \frac{4\Delta t}{2\Delta h_j} - u_{j-2}^{n+1} g_{j-2}^n \frac{1\Delta t}{2\Delta h_j}}{1 + \mu_j^n \Delta t + g_j^n \frac{3\Delta t}{2\Delta h_j}},\\
B &= s_{\mathrm{e}} \sum_{j=0}^{J-1} \beta_j^n u_j^n h_j.
\end{aligned} \tag{S6}
$$

where we have used the first-order scheme to calculate $u_1$.

#### Explicit escalator-boxcar-train method (EBT)

The EBT scheme divides the population into cohorts, and then tracks the mean body size and abundance of individuals within each cohort. Here, we present the EBT equations as implemented in libpspm for completeness, and refer the reader to (Brännström et al., 2013; de Roos, 1988) for further details on the EBT scheme. In our implementation, cohorts are sorted from largest to smallest by size. Thus, the boundary cohort is at the index $J-1$, whereas internal cohorts are indexed by $i \in [0,1,2,\ldots,J-2]$. The dynamics of internal cohorts are described by

$$
\begin{aligned}
\frac{\mathrm{d}N_i}{\mathrm{d}t} &= -\mu(x_i,t,E)N_i,\\
\frac{\mathrm{d}x_i}{\mathrm{d}t} &= g(x_i,t,E).
\end{aligned} \tag{S7}
$$

The dynamics of the boundary cohort are described by tracking the cumulative size increment $\pi_{\mathrm{b}}$ of offspring over their birth size, rather than the absolute size,

$$
\begin{aligned}
\frac{\mathrm{d}N_{\mathrm{b}}}{\mathrm{d}t} &= -\mu(x_{\mathrm{b}},t,E)N_{\mathrm{b}} - \mu_x(x_{\mathrm{b}},t,E)\pi_{\mathrm{b}} + B,\\
\frac{\mathrm{d}\pi_{\mathrm{b}}}{\mathrm{d}t} &= g(x_{\mathrm{b}},t,E)N_{\mathrm{b}} + g_x(x_{\mathrm{b}},t,E)\pi_{\mathrm{b}} - \mu(x_{\mathrm{b}},t,E)\pi_{\mathrm{b}},\\
x_{J-1} &= x_{\mathrm{b}} + \frac{\pi_{\mathrm{b}}}{N_{\mathrm{b}}},\\
N_{J-1} &= N_{\mathrm{b}},\\
B &= s_{\mathrm{e}} \sum_{i=0}^{J-1} \beta(x_i,t,E)N_i.
\end{aligned} \tag{S8}
$$

The gradients $g_x$ and $\mu_x$ required in Eq. (S8 are computed numerically. For the explicit method, we integrate the derivatives in Eq. (S8 using the Runge-Kutta 4-5 Cash-Karp integrator.

Periodically, after a time interval $\Delta T_{\mathrm{b}}$, the boundary cohort is internalized, and a new boundary cohort is inserted into the population. When the boundary cohort is internalized, the values of $x_{J-1}$ and $N_{J-1}$ are set according to Eq. (S8. Cohorts in which the number of individuals have fallen below a threshold value $u_{\mathrm{cut}}$ are removed from the population. Normally, when cohorts are numbered in ascending order, insertion of a new boundary cohort requires a renumbering so that the new boundary cohort gets the index 0, but this is not necessary in libpspm as cohorts are sorted descending. Rather only the total number of cohorts $J$ needs to be incremented by 1, so that indices of all cohorts remain the same, and the new boundary cohort gets the new index $J-1$ (with incremented $J$).

#### Semi-implicit escalator-boxcar-train method (IEBT)

For the semi-implicit method, we use a semi-implicit time update in Eq. (S2 and Eq. (S8, to get

$$\begin{aligned}
\frac{N_i^{n+1}-N_i^n}{\Delta t} &= -\mu_i^n N_i^{n+1}, \\
\frac{x_i^{n+1}-x_i^n}{\Delta t} &= g_i^n, \\
\frac{N_\mathrm{b}^{n+1}-N_\mathrm{b}^n}{\Delta t} &= -\mu_\mathrm{b}^n N_\mathrm{b}^{n+1} - \mu_{\mathrm{b}x}^n \pi_\mathrm{b}^{n+1} + B, \\
\frac{\pi_\mathrm{b}^{n+1}-\pi_\mathrm{b}^n}{\Delta t} &= g_\mathrm{b}^n N_\mathrm{b}^{n+1} + g_{\mathrm{b}x}^n \pi_\mathrm{b}^{n+1} - \mu_\mathrm{b}^n \pi_\mathrm{b}^{n+1}, \\
B &= s_\mathrm{e} \sum_{i=0}^{J-1} \beta_i^n N_i^n.
\end{aligned} \tag{S9}$$

The first two equations in this system can be solved to give the values of $N_i^{n+1}$ and $x_i^{n+1}$,

$$\begin{aligned}
N_i^{n+1} &= N_i^n/(1+\mu_i^n \Delta t), \\
x_i^{n+1} &= x_i^n + g(x_i, t, E)\Delta t.
\end{aligned} \tag{S10}$$

The last two equations form a system of two simultaneous equations for $N_\mathrm{b}^{n+1}$ and $\pi_\mathrm{b}^{n+1}$, as follows,

$$\begin{bmatrix} 1+\mu_\mathrm{b}^n \Delta t & \mu_{\mathrm{b}x}^n \Delta t \\ -g_\mathrm{b}^n \Delta t & 1 - g_{\mathrm{b}x}^n \Delta t + \mu_\mathrm{b}^n \Delta t \end{bmatrix} \begin{bmatrix} N_\mathrm{b}^{n+1} \\ \pi_\mathrm{b}^{n+1} \end{bmatrix} = \begin{bmatrix} N_{\boldsymbol{i}}^n + B\Delta t \\ \pi_\mathrm{b}^n \end{bmatrix}, \tag{S11}$$

which can readily be solved for $N_\mathrm{b}^{n+1}$ and $\pi_\mathrm{b}^{n+1}$.

#### Characteristic method (CM)

The characteristic method solves the PDE in Eq. 1 by finding “characteristic curves” along which the PDE reduces to an ordinary differential equation (ODE), and then integrating the resulting ODEs using the Runge-Kutta 4-5 Cash-Karp integrator. It can be shown that each characteristic curve can be found by solving the ODE

$$\frac{\mathrm{d}x_i(t)}{\mathrm{d}t} = g(x_i, t, E),$$

$$x_i(0) = x_\mathrm{b},$$

where $t$ is the time since the insertion of the characteristic curve. The density function along the characteristic curve is given by the solution of

$$\frac{\mathrm{d}u(x_i,t)}{\mathrm{d}t} = -g_x(x_i,t,E)u(x_i,t) - \mu(x_i,t,E)u(x_i,t).$$

The initial density of characteristic curves is set at the time of insertion of the curve into the system. After each time interval $\Delta T_\mathrm{b}$, a new characteristic curve is inserted into the system, and characteristic curves whose distance to neighbouring curves is less than a threshold $x_{\text{cut}}$, or whose density is less than a threshold $u_{\text{cut}}$, are removed from the population. The initial density of a new curve, being inserted at time $t_0$ with index $J-1$ (similar to the numbering in the EBT scheme) is calculated using the boundary condition,

$$u(x_{J-1},0)=\frac{s_{\text{e}}}{g(x_{\text{b}},0,E(t_0))}\sum_{i=1}^{J-2}\left(\frac{u_{i-1}\beta_{i-1}(t_0)+u_i\beta_i(t_0)}{2}\right)(x_i-x_{i-1}).$$

Note that we use trapezoidal integration to calculate the total flux of offspring.

#### Semi-implicit characteristic method (ICM)

In the semi-implicit version of the characteristic method, $u(x_i,t+\Delta t)$ is given by

$$u(x_i,t+\Delta t)=\frac{u(x_i,t)}{1+g_x(x_i,t,E)\Delta t+\mu(x_i,t,E)\Delta t}.$$

#### Agent-based method (ABM)

The agent-based method does not involve solving any differential equations. Instead, we simulate a large population of $J$ agents, with each agent $i$ representing a fixed number $N_{\text{s}}$ of identical real individuals. At each time step of length $\Delta t$, the population goes through the following processes, in this order.

*Reproduction* – All individuals reproduce to give the total number of offspring according to the following equation, rounded to the nearest integer number of agents,

$$B=s_{\text{e}}\sum_{i=0}^{J-1}\beta_i^n N_{\text{s}}.$$

*Mortality* – Individuals are removed from the population with an independent probability of dying in the time interval $\Delta t$, given by

$$p_{\text{die}}=1-e^{-\mu_i^n\Delta t}.$$

*Growth* – Surviving individuals grow in size according to the equation

$$x_i^{n+1}=x_i^n+g_i^n\Delta t.$$

*Establishment* – The offspring calculated in the first step are inserted into the population, creating an equivalent ($B/N_{\text{s}}$) number of new agents.

### Supplementary figures

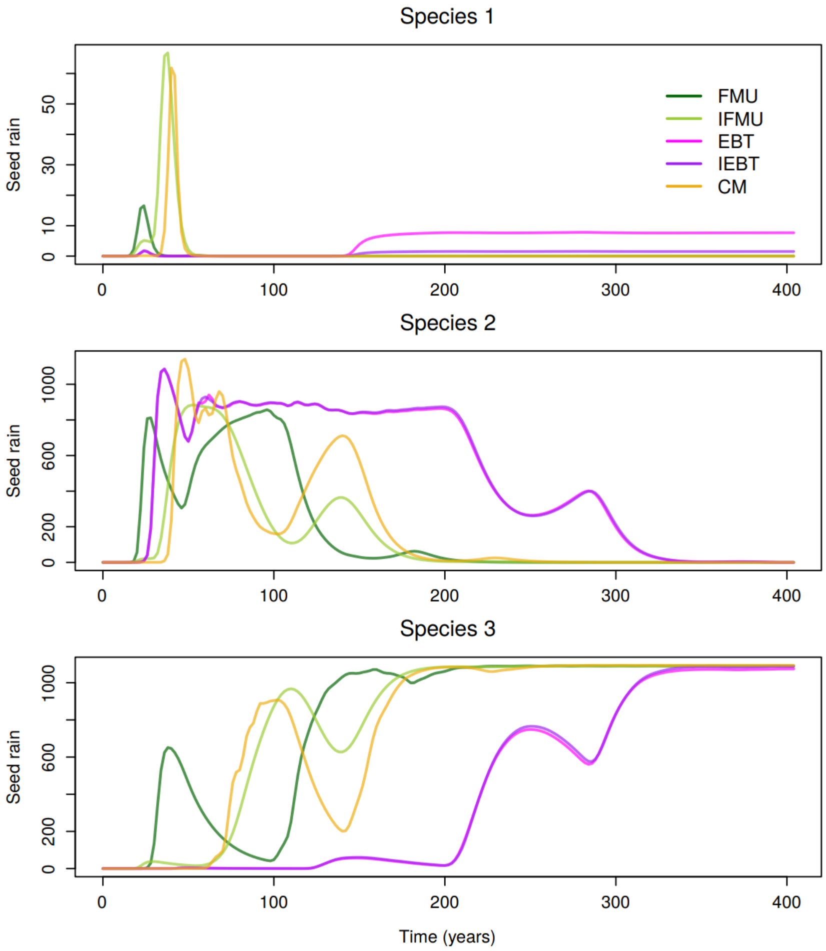

**Fig. S1. Transient solutions of the three-species Plant model.** The dynamics are simulated in feedback mode with the initial condition $u(x, 0) = u_0\delta(x - x_{\mathrm{b}})$, where $\delta(x)$ is the Dirac delta function. The eight methods disagree on the solution.

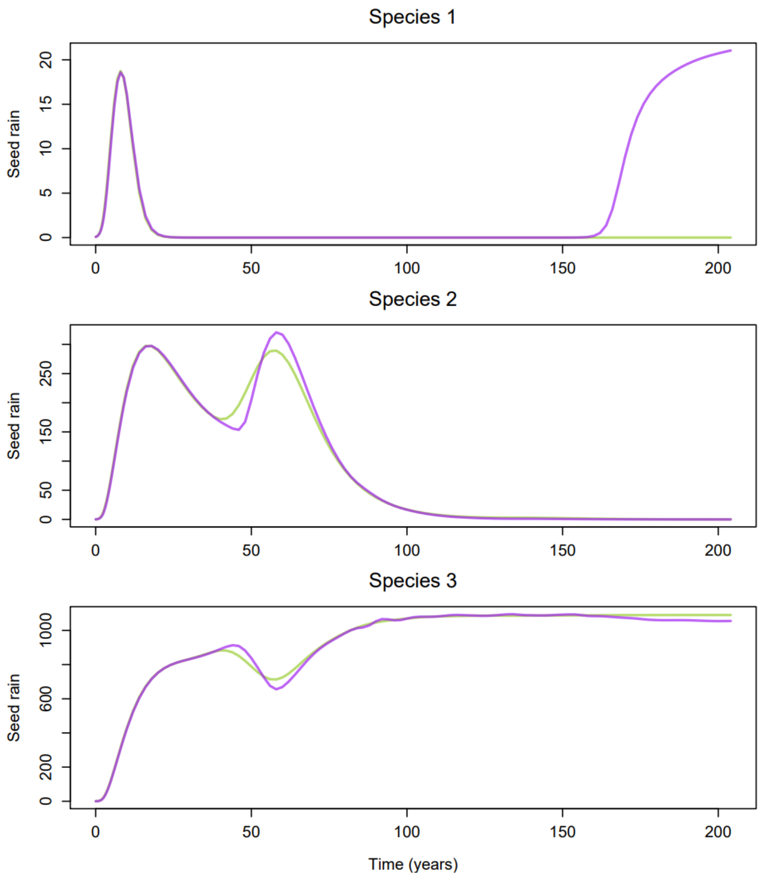

**Fig. S2. Transient solutions of the three-species Plant model.** The dynamics are simulated in feedback mode with the initial condition $u(x,0) = u_0 e^{-x/4}$. The IEBT (purple) and IFMU (light green) are the only methods that can successfully simulate this scenario.

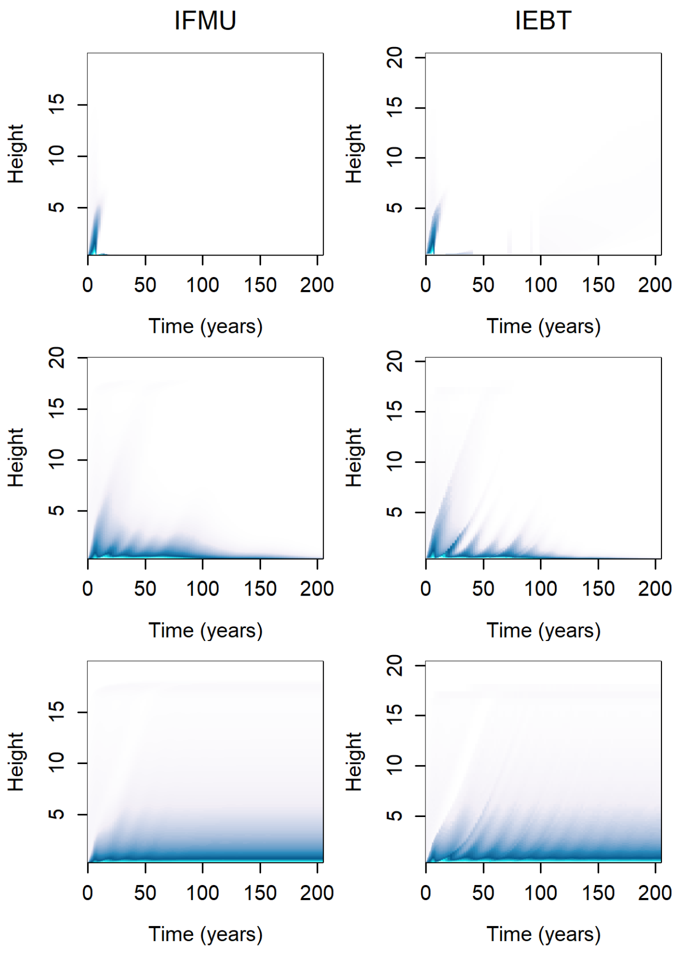

**Fig. S3. Size distributions of the three-species Plant model.** The dynamics are simulated in feedback mode with the initial condition $u(x, 0) = u_0 e^{-x/4}$ using the IFMU and IEBT methods.
